# Optogenetic Protein Cleavage in Zebrafish Embryos

**DOI:** 10.1101/2022.05.23.493165

**Authors:** Wes Brown, Savannah Albright, Michael Tsang, Alexander Deiters

## Abstract

A wide array of optogenetic tools is available that allow for precise spatiotemporal control over many cellular processes. These tools have been especially popular among zebrafish researchers who take advantage of the embryo’s transparency. However, photocleavable optogenetic proteins have not been utilized in zebrafish. We demonstrate successful optical control of protein cleavage in embryos using PhoCl, a photocleavable fluorescent protein. This optogenetic tool offers temporal and spatial control over protein cleavage events, which we demonstrate in light-triggered protein translocation and apoptosis.

## Introduction

Zebrafish embryos are an excellent vertebrate model system used across many fields from developmental biology and neuroscience, to drug discovery and the biology of aging.^1,2^ Outside of their ex utero development, small size, ease of genetic manipulation, and expression of injected mRNA, one of the main advantages of working with zebrafish is the transparency of the embryo and larva, allowing for visualization of organ development and fluorescent imaging in live animals.^3^ Their transparency also makes them amenable to using light as an control tool that offers high spatial and temporal precision. Optogenetics has garnered many successful uses in zebrafish embryos, including control over gene expression,^4^ cell migration,^5^ protein localization,^6^ cell signaling,^7^ and cell ablation.^8^ These tools have been used to study important biological processes due to the unparalleled spatiotemporal control afforded by light.^9^ Unlike optogenetically regulated dimerization or conformational changes of proteins, optically induced bond cleavage provides permanent activation or deactivation of protein activity without the need for continuous illumination. In zebrafish embryos, we have successfully photo-controlled kinase^10^ and Cre recombinase^11^ activity through genetic code expansion with amino acids carrying photocleavable caging groups on their sidechains.^19–20^ Optical control of Cas9 gene editing^12–15^ and base editing^16^ using a guide RNA containing photocaged nucleobases has been established. Additionally, control of gene expression with caged morpholino oligonucleotides by circularizing with a photocleavable linker or through photocaged nucleobases has been reported.^17–20^

In seminal work, the Campbell lab has engineered the green to red photoconvertible fluorescent protein mMaple to cleave into a 230 residue N-terminal and 12 residue C-terminal fragments after violet light illumination (**Figure 1A**).^21^ When the green fluorescent chromophore is irradiated by violet light, it undergoes covalent bond breakage at the protein backbone while also destroying the chromophore (**Figure 1B**). The engineered PhoCl construct was used in mammalian cells to remove a NES or NLS from a fluorescent protein, activate Gal4 or Cre recombinase activity by optically cleaving off steroid receptors, and activate a protease by removing an inhibitory peptide. More recently, they have further optimized their photocleavable protein to improve fragment dissociation kinetics and yield, labeling the newly optimized construct PhoCl2c.^22^ Another variant, PhoCl2f, was also generated and had comparable dissociation kinetics as PhoCl2c, but PhoCl2c had better overall activation of a nuclear translocation reporter, making it the favored candidate. PhoCl has since been used by other labs to generate tunable hydrogels,^23^ release proteins from biomaterials,^24, 25^ study cell mechanotransduction,^26^ and generate membraneless organelles.^27^ PhoCl has similar benefits as other light-induced bond cleavage systems described above, namely allowing for rational design of the light-activated system and not requiring continuous illumination for sustained activity, which is advantageous for precise control in a rapidly developing, multicellular organism, but it also shares the accessibility and ease of use of a fully genetically encoded optogenetic tool. Heterologous expression and chromophore development is efficient, providing for robust activation of protein function.

**Figure 1.**
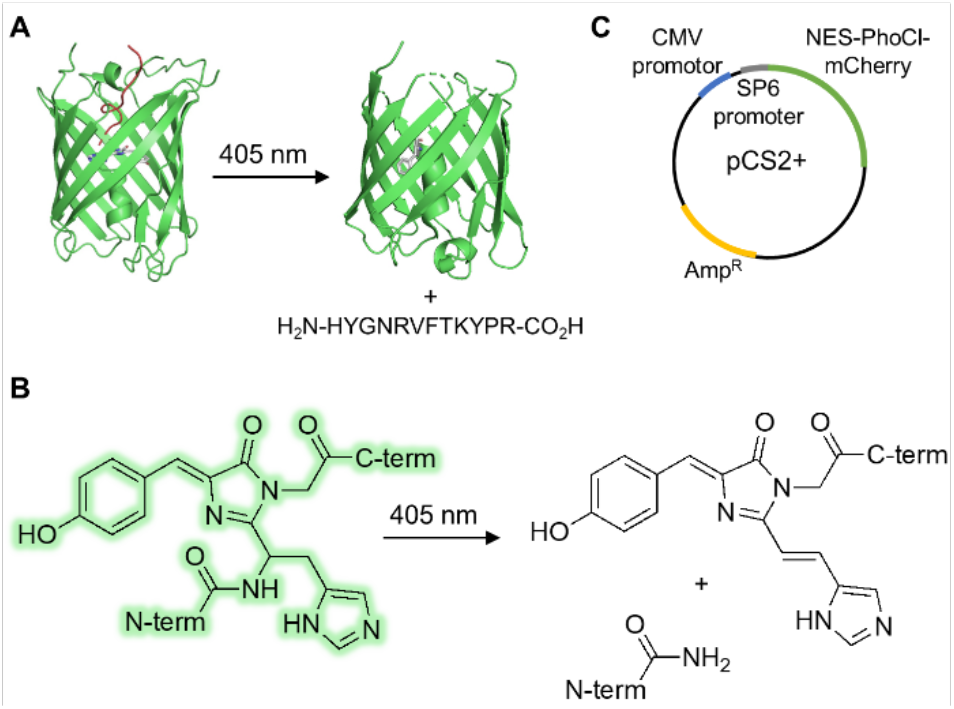
A) PhoCl structure before and after photocleavage. B) Mechanism of PhoCl backbone cleavage after irradiation of the chromophore. C) pCS2 vector used for both mammalian cell and zebrafish embryo experiments.

## Results and Discussion

We chose the optimized PhoCl2c for our studies in embryos (referred to as just PhoCl here), specifically optical control of nuclear exclusion of a fluorescent reporter (NES-PhoCl-mCherry) and optical control of induction of apoptosis (NBid-PhoCl-CBid).^22^ The corresponding genes were synthesized with human-optimized codon usage since difference between both species are minimal and cloned into a pCS2+ vector via Gibson assembly (**SI Table 1-2**). This vector contains a CMV promotor for mammalian expression as well as an Sp6 promotor for mRNA generation for zebrafish injections, making it a versatile tool for use in both cell culture and embryos (**Figure 1C**).

Photocleavage of NES-PhoCl-mCherry results in removal of a nuclear exclusion sequence (NES) from the fused mCherry and subsequent diffusion of the fluorescent reporter from the cytoplasm into the nucleus (**Figure 2A**). Nuclear exclusion can be quantified by determining the nuclear to cytoplasmic (N/C) mCherry fluorescence intensity ratio of single cells in micrographs. Diffusion of mCherry to the nucleus produces a consistent red fluorescent intensity throughout the cell and therefore should result in an ideal N/C ratio of 1. To validate the function of the NES-PhoCl-mCherry construct, HeLa cells expressing it were irradiated at 405 nm for 5 minutes, ensuring complete photoconversion. The cells irradiated with 405 nm light showed immediate loss of green fluorescence, indicating PhoCl photocleavage (**Figure 2B**). mCherry localization was tracked for 35 minutes through time-lapse imaging and the nuclear to cytoplasmic ratio of mCherry fluorescence intensity for single cells was determined. Briefly, ImageJ was used to measure the mean gray value of a region within the nucleus and a region in the cytoplasm. Background was subtracted and values were divided for the N/C ratio. This was repeated for each cell and averaged for each timepoint. Before irradiation, nuclear exclusion was observed, followed by rapid movement of mCherry into the nucleus visible at 2 minutes. By 35 minutes, the average N/C fluorescence in the irradiated cells was 0.93, suggesting virtually complete redistribution of mCherry (**Figure 2C**). Therefore, PhoCl offers the ability to directly manipulate protein translocation through optical cleavage of localization signals.

**Figure 2.**
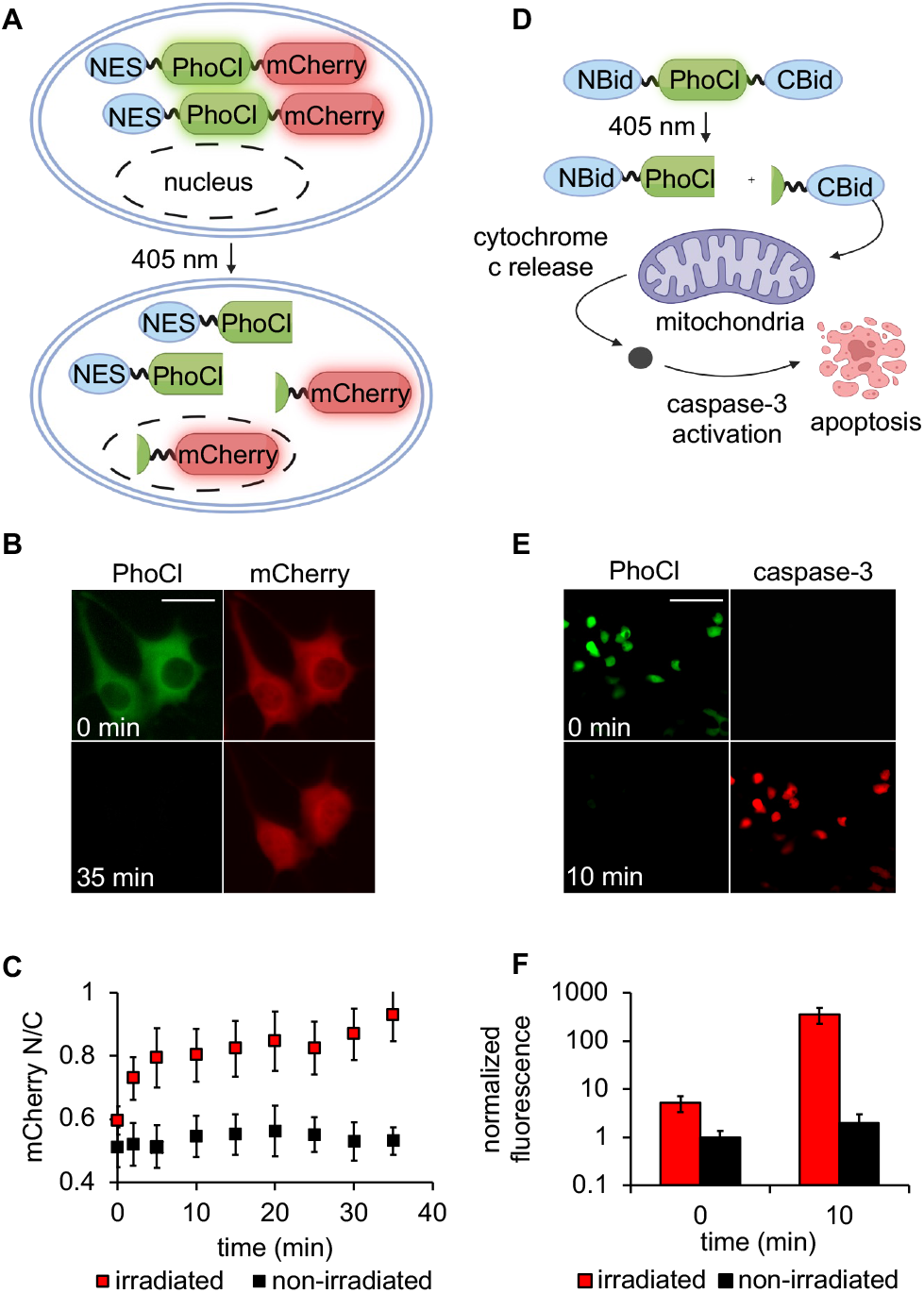
Photoactivation of PhoCl in mammalian cells. A) NES-PhoCl-mCherry photoconversion and dissociation. B) Nuclear translocation of mCherry tracked for 35 minutes post 405 nm irradiation of HeLa cells expressing NES-PhoCl-mCherry. Scale bar = 20 μm. C) Nuclear to cytoplasmic mCherry fluorescence ratio was calculated for irradiated and non-irradiated cells and plotted over time. Each point represents the mean from 10 cells and error bars represent standard deviations. D) NBid-PhoCl-CBid photoconversion and dissociation leading to apoptosis of irradiated cells. E) Caspase activation tracked for 10 minutes post 405 nm irradiation of HEK293T cells expressing NBid-PhoCl-CBid. Scale bar = 100 μm. F) Caspase-3 activity (NucView 530 fluorescence) was measured for irradiated and non-irradiated cells. Each bar is the mean of 9 fields of view, normalized to background fluorescence of non-irradiated cells at 0 min. Error bars represent standard deviations.

We next wanted to validate another construct exhibiting control over protein function through PhoCl photocleavage for later use in zebrafish embryos. Bid is a proapoptotic cytoplasmic protein that when cleaved by caspase-8 into the active, truncated form tBid activates apoptosis through translocation to the mitochondria, inducing an increase of membrane permeability and cytochrome c release (**Figure 2D**).^24^ By replacing the caspase-8 cut site with PhoCl, spatiotemporal control of Bid cleavage, and therefore the apoptosis pathway, can be achieved.^11^ HEK293T cells transiently expressing NBid-PhoCl-CBid after overnight transfection were analyzed for the ability to optically trigger caspase activity. The NucView 530 caspase-3 reporter (Biotium) is a fluorogenic dye that is cell permeable and upon cleavage of its caspase-3/7 recognition peptide DEVD, binds to DNA thereby becoming fluorescent. The reporter was added to cell media at a final concentration of 2 μM and cells were irradiated with a 405 nm LED and imaged over 40 minutes. Irradiated cells exhibited a loss of green fluorescence, due to PhoCl cleavage, and significant red fluorescence at 10 minutes, due to rapid caspase activation, compared to cells not exposed to light (**Figure 2E**). Photostimulation led to a 300-fold increase in caspase activity compared to cells kept in the dark and remained active for 40 minutes, as determined by intensity measurements in the red fluorescence channel (**Figure 2F, SI Figure 1**). Confirmation of expression and optogenetic function of the pCS2-PhoCl constructs in mammalian cells set the stage for applications in zebrafish embryos. To date, photocleavable proteins have not been used in zebrafish embryos. First, we generated the mRNA encoding the NES-PhoCl-mCherry reporter through an *in vitro* transcription of the pCS2 plasmid (**SI Figure 2**) and injected it into zebrafish embryos at the one-cell stage (**Figure 3A**). At 24 hour postfertilization, the whole embryo was irradiated with a 405 nm LED, and the animals were anesthetized and immobilized in low-melting point agarose for confocal imaging 40 minutes after light exposure. Embryos kept in the dark had green fluorescence and saw complete nuclear exclusion of mCherry, while irradiated embryos showed loss of green fluorescence and mCherry was equally distributed throughout the cell as expected (**Figure 3B**). The loss of green fluorescence is unique to PhoCl compared to EGFP-mCherry protein that was stable under the same irradiation conditions (**SI Figure 3**). Thus, the loss of green fluorescence can be utilized as an indicator of successful PhoCl cleavage and reveals the extent and localization of photoactivation, which is particularly useful for spatially restricted irradiation of model organisms.

**Figure 3.**
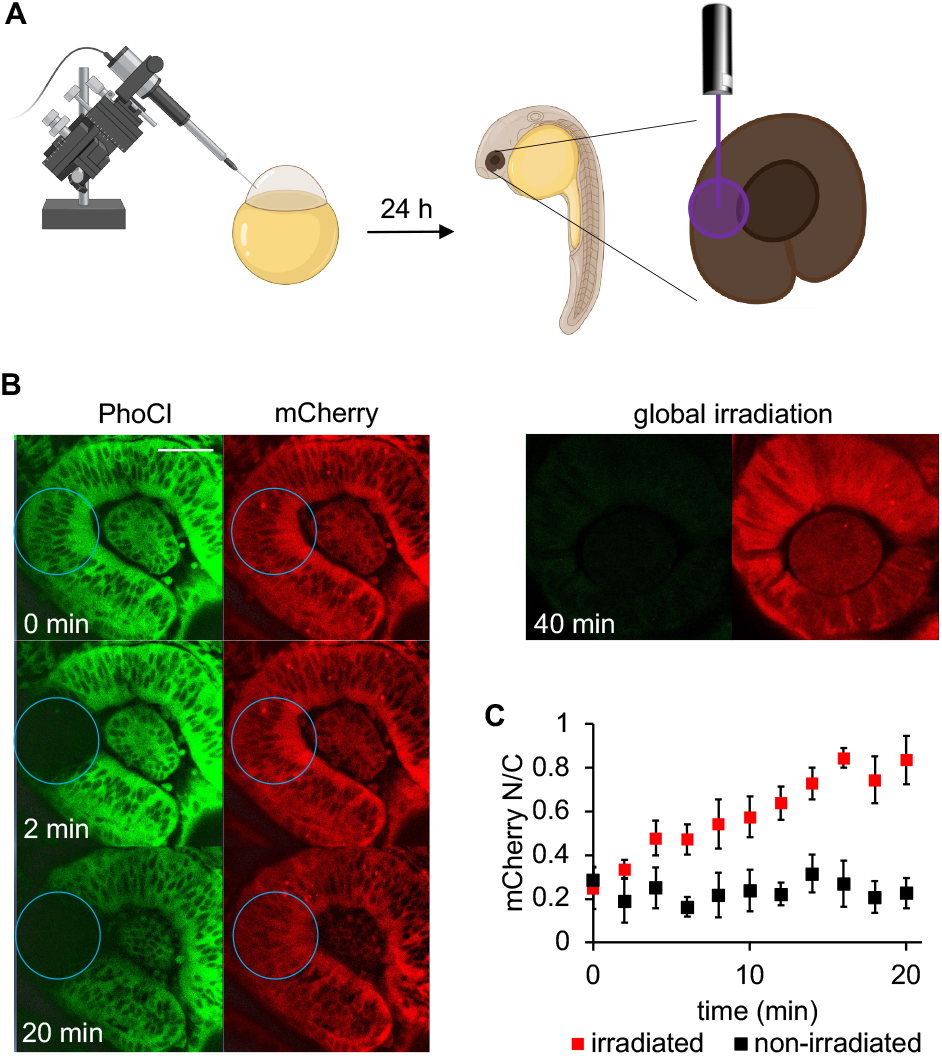
Optical control of nuclear translocation using PhoCl in zebrafish embryos. A) Embryos at 24 hpf expressing NES-PhoCl-mCherry subjected to targeted irradiation (blue circle) of the retina. Nuclear translocation of mCherry was tracked over 25 minutes. B) An embryo at 24 hpf expressing NES-PhoCl-mCherry was subjected to global irradiation and imaged at 40 min post irradiation. Timelapse images of localized activation of nuclear translocation is shown on the left. C) Nuclear to cytoplasmic ratio was calculated for irradiated and non-irradiated cells within the same images and plotted over time. Each point represents the mean of 5 cells and error bars represent standard deviations. Scale bar = 40 μm.

Optical activation enables precise spatial control over the protein nuclear translocation. A small circular area was irradiated within the embryo eye by using the bleach function to irradiate with the 405 nm laser for 20 iterations at 30% laser power and a localized loss of green fluorescence was visible immediately. The irradiated cells were tracked for 20 minutes and a loss of nuclear exclusion was observed only within the irradiated area (**Figure 3B, SI Figure 4**). The nuclear to cytoplasm ratio of mCherry fluorescence was measured over time by tracking mCherry fluorescent intensity in single cells. A significant increase in nuclear fluorescence was observed in the irradiated cells compared to the non-irradiated cells and was complete by 16 minutes (**Figure 3C**), matching the results seen in cell culture experiments (**Figure 2C**). Thus, PhoCl photocleavage is a viable option for acutely removing protein localization sequences and triggering protein translocation in zebrafish embryos. This allows for mislocalization of a protein of interest and activation of protein function through translocation to its native cellular compartment, all with high spatiotemporal resolution in a live animal.

Having successfully controlled protein localization with PhoCl in embryos, we endeavored to activate apoptosis with spatiotemporal resolution in embryos as well. mRNA was made for NBid-PhoCl-CBid (**SI Figure 2**) and injected into one-cell stage embryos. Protein was expressed until 24 hpf, showing strong green fluorescence of the photocleavable system. The same peptidic caspase-3 reporter (NucView 530) was added to the fish water^28^ and embryos were irradiated with a 405 nm LED and imaged over 40 minutes (**Figure 4A**).^25^ Red fluorescence was observed by 10 minutes in the irradiated embryo, and by 20 minutes tail curling and yolk lysis can be seen, indicating caspase activation and subsequent embryo apoptosis, while simultaneous irradiation conditions in non-injected embryos induced no caspase activation (**SI Figure 5**). By 40 minutes, nearly all embryos had fully lysed. (**SI Figure 6**, **SI Video 1,** N = 27). Tail curling and cell lysis have been observed with activation of apoptosis previously.^29, 30^ The dramatic lethality of cleaved NBid-PhoCl-CBid is expected due to the acute, large pulse of cleaved Bid that plays a key role in both ligand-activated extrinsic and cytotoxic stress-induced intrinsic apoptosis. Next, we performed spatially restricted activation of apoptosis through localized irradiation near the epidermis of the embryo tail, showing red fluorescence within 2 minutes as well as apoptosis of the irradiated cells, which progressed to an epidermal lesion (**Figure 4B**). Irradiation was also performed in the embryo eye and red fluorescence was seen within 2 minutes limited only to the irradiated cells (**SI Figure 7**). This method of apoptosis induction and cell ablation is particularly exciting due to the precise optical control and rapid onset. Other systems for light induced cell ablation are based on optogenetically controlled transcriptional activation of nitroreductase with subsequent metronidazole addition, or mutant M2 proton channel expression originally from influenza.^4^ The NBid-PhoCl-CBid system induces immediate caspase cascade activation and is easy to use since it only requires injection of a single mRNA.

**Figure 4.**
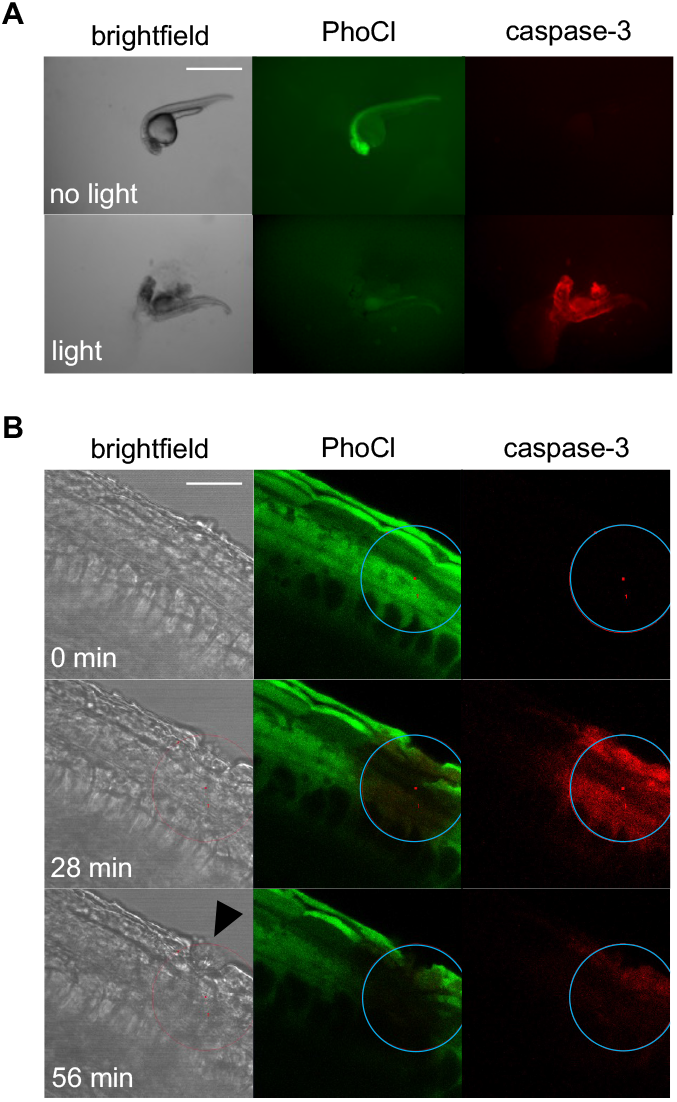
Control of caspase-3 activation using PhoCl in zebrafish embryos. A) Activation of a fluorescent caspase reporter through whole embryo irradiation of animals expressing NBid-PhoCl-CBid. Scale bar = 1.5 mm. B) Targeted irradiation (blue circle) of the tail epidermis at 24 hpf and imaging over 56 minutes. The black arrow indicates the epidermal defect seen after irradiation. Scale bar = 40 μm.

## Conclusion

In conclusion, light-induced protein backbone-cleavage is a powerful tool that is complemented by the optical transparency of the zebrafish embryo, a well-established model organism for human physiology and disease. Additional advantage of zebrafish include their low cost of maintenance, vertebrate biology with high genetic similarity to humans, and ease of protein expression and genetic manipulation.^31–33^ We assembled convenient expression constructs that are functional in both mammalian and zebrafish systems to demonstrate optical control over nuclear exclusion and apoptosis, which can be applied to control protein compartmentalization and ablate cells with high spatiotemporal resolution. Photocleavage was induced with 405 nm light, making it amenable to experiments on most epifluorescence and point-scanning confocal microscopes, and physiological responses to our reporter activation occurred within minutes, suggesting rapid dissociation of the cleaved PhoCl fragments, similar to mammalian cell experiments.^22^ PhoCl can easily be applied to control protein localization via cleavage of NES, NLS, or membrane anchors, thereby enabling temporary protein mislocalization into a particular compartment as an approach for conditional control. The NBid-PhoCl-CBid construct induces rapid apoptosis and cell ablation, which can be utilized in studies of tissue response to damage and the developmental impacts of certain cell populations. This method of cellular ablation has advantages compared to other established laser based methods,^34–36^ namely requiring minimal 405 nm laser exposure, thereby preventing injury and stress response of neighboring cells, simple and effective design, and no need for specialized equipment or reagents outside of a standard microscope and an *in vitro* transcription kit. This optogenetic system may also have further utility through the generation of a transgenic fish line thereby enabling applications in adult animals. To our knowledge, this is the first demonstration of a photocleavable protein in zebrafish embryos. We expect PhoCl to be a very accessible and broadly applicable optogenetic tool for zebrafish researchers across many fields since, as we have demonstrated here, PhoCl constructs developed in mammalian cells can be readily translated to the zebrafish model.

## Experimental Section

### Cell culture maintenance and transfection

All cell culture experiments were performed in a sterile laminar flow hood. HeLa and HEK 293T cells were maintained in Dulbecco’s Modified Eagle Medium (DMEM, Gibco) supplemented with 10% (v/v) fetal bovine serum (Sigma-Aldrich) and 1% (v/v) penicillin/streptomycin at 37 °C with 5% CO2. Cells were used between passage number 6 and 25 and were tested for mycoplasma contamination every 6 months. Cells were seeded in a 96 well μClear plate and in 100 μL of DMEM (antibiotic free).

For translocation assays, when HeLa cells reached 80-90% confluency, each well was transfected with 200 ng of plasmid using 0.4 μL of P3000 and 0.4 μL of Lipo3000 (ThermoFisher), diluted in Opti-MEM transfection media (Thermo Scientific) to a total volume of 10 μL per well according to manufacturer instructions. For caspase activation, when HEK293T cells reached 80-90% confluency, each well was transfected with 100 ng of plasmid using 0.4 μL of P3000 and 0.4 μL of Lipo 3000, diluted in Opti-MEM transfection media to a total volume of 10 μL per well according to manufacturer instructions. Cells were incubated at 37 °C with 5% CO2 for 24 hours before imaging.

### Mammalian cell irradiation and imaging

Cell irradiation (5 minutes) was performed using a 405 nm LED (Luxeonstar, Luxeon Z, 675 mW), placed 6 cm above the 96 well plate containing the cells suspended in 40 μL Fluorobrite media. Light output at the specimen was measured at 80 mW with a Thorlabs power meter. Cells were imaged using a Zeiss Axio Observer Z1 with an Andor Zyla 4.2 camera with a GFP filter (ex: 430-510, em: 475-575), a DsRed filter (ex. 525-575, em: 635-675), and the brightfield channel. For caspase-3 activation experiments, NucView 530 (Biotium) was included in Fluorobrite at a final concentration of 2 μM for visualization of caspase-3 activity in the DsRed channel. N/C ratios were quantified in ImageJ by measuring the mean fluorescence intensity of a representative area within the cellular regions of interest and averaging individual N/C values. Standard error was determined from 10 individual cells combined from three biological replicates. Caspase activation images were analyzed in ImageJ. A threshold was generated using PhoCl fluorescence (490 nm) to remove background and select fluorescent cells. The selection was applied to the corresponding NucView 530 channel (530 nm) and the mean fluorescence intensity was measured at each timepoint. Standard error was determined from averaging 9 fields of view.

### Zebrafish care and microinjection

The zebrafish experiments were performed according to a protocol approved by the Institutional Animal Care and Use Committee (IACUC) at the University of Pittsburgh (protocol no 19075360). Embryos were collected after natural mating of the AB* fish line. Injection solutions of the NES-PhoCl-mCherry or NBid-PhoCl-CBid mRNAs were prepared at 400 or 200 ng/μl, respectively, with phenol red added to a final concentration of 0.05% as a tracer for injection. A volume of 2 nL was injected into the yolk of 1-2 cell stage embryos using a World Precision Instruments Pneumatic PicoPump injector. Embryos were incubated in E3 water at 28.5 °C for 24 hours in the dark before imaging.

### Zebrafish irradiation and imaging

For whole embryo irradiation (5 minutes), a 405 nm LED (Luxeonstar, Luxeon Z, 675 mW) was placed 3 cm above the 35 mm petri dish containing the embryos suspended in E3 water. Light output at the specimen was measured at 350 mW with a Thorlabs power meter. Stereoscope imaging of embryos was performed with a Leica M205 FA microscope with a DsRed filter (ex: 510-560, em: 590-650), an EGFP filter (ex: 450-490 nm, em: 500-550 nm), and the bright field channel. A Zeiss LSM 700 laser scanning confocal microscope was used for spatial activation experiments. At 24 hpf the embryos were embedded in 500 μl of 1.5 % low-melting-point agarose in a glass bottom 35 mM dish. The bleach function was used with the 405 nm laser (5 mW) at 30% laser power and 20 iteration and a pixel dwell time of 3.2 μs. The regions function was used to limit irradiation to selected cell populations. After irradiation, time lapse imaging was performed every 2 minutes. The N/C ratio was calculated using ImageJ software by measuring background subtracted mean grey value for the nuclear and cytoplasmic region for five cells. For caspase-3 activation experiments, NucView 530 (Biotium) was included in the embryo water at a final concentration of 2 μM for visualization of caspase-3 activity with the DsRed filter on the Leica stereoscope or 555 nm laser of the Zeiss confocal microscope. PhoCl was imaged with the EGFP filter described above. For whole-embryo time-course caspase activation experiments, images were taken every minute for 40 min and the video was rendered in Adobe Photoshop.

## Acknowledgements

We acknowledge financial support from the National Science Foundation (CHE-1904972). WB was supported by a University of Pittsburgh Mellon Fellowship. We would also like to thank Dr. Juhoon So for assistance with confocal imaging of zebrafish embryos. Some figures were generated using BioRender.com.

