## Supporting Information for "Optogenetic Protein Cleavage in Zebrafish Embryos"

### Additional Methods

#### Plasmid construction

Both the NES-PhoCl-mCherry and NBid-PhoCl-CBid constructs were cloned into pCS2+. The pCS2+ backbone (4 µg) was digested with Bamh1 and EcoR1 (NEB) in a 10 µl reaction for 1 hour at 37 °C. It was loaded onto a 0.8% agarose gel and electrophoresed, and the vector product was excised for in gel extraction and purification using the GeneJet gel purification kit (Thermo) according to the manufacturer's protocol. The NES-PhoCl and NBid-PhoCl-CBid gene sequences were obtained from Addgene (#164037 and #164051) and gene fragments were ordered (Twist Biosciences). Primers 1 and 2 (NES-PhoCl) or 3 and 4 (mCherry) or 5 and 6 (NBid-PhoCl-CBid) were used to amplify the gene products in 50 µl PCR reactions. Vector overlaps of 30 bp were included for Gibson assembly later. The reactions were loaded on a 0.8% agarose gel and electrophoresed, and correct sized bands were excised and purified as mentioned above. Next, the purified gene products were used in a 20 µl Gibson assembly reactions at 50 °C for 1 hour with 100 µg of pCS2+ digested vector and 2 molar equivalents of the insert fragments. Top 10 chemically competent cells (50 µl) were transformed with 5 µl of the Gibson assembly reaction, plated on ampicillin containing LB agar, and colonies were selected for growth in LB media containing ampicillin after overnight incubation at 37 °C. Overnight cultures in LB media were miniprepped using the GeneJet plasmid miniprep kit (Thermo) following manufacturer's protocol to yield pCS2+-NES-PhoCl-mCherry and pCS2+-NBid-PhoCl-CBid.

#### In vitro transcription

To make mRNA, 4 µg of plasmid was linearized with Not1 (NEB) in a 10 µl reaction at 37 °C for 1 hour. It was diluted with water to a total volume of 50 µl and purified by phenol: chloroform: isoamyl alcohol (50 µl) extraction, precipitated in 138 µl of ethanol at -20 °C overnight, then pelleted at 15,000 rpm for 5 minutes. The DNA pellet was washed with 70% ethanol, repelleted, the supernatant was removed, and then the pellet was dried at room temp with the cap open before resuspension in 20 µl RNase free water. The linearized plasmid (1 µg) was used in a 20 µl Sp6 mMessage mMachine in vitro transcription reaction following manufacturer instructions. Reaction mixtures were treated with 1 µl of Turbo Dnase (Thermo), purified by phenol: chloroform: isoamyl alcohol extraction, ethanol precipitated, and resuspended in Rnase free water as described above and stored at -80 °C.

### Supporting Tables

| Primer | Sequence (5' to 3') |
|---|---|
| 1 | TACAAGCTACTTGTTCTTTTTGCAGGATCCATGAACCTGGTGGACCTGCAGAAGAAGC |
| 2 | GCTGCCCCCGCCTCCGCTGCCCCCGCCTCCGGTACCCCGTGGGTACTTGGTGAACACGCG |
| 3 | GGTACCGGAGGCGGGGGCAGCGGAGGCGGGGGCAGCGTGAGCAAGGGCGAGGAGGATAAC |
| 4 | AGTTCTAGAGGCTCGAGAGGCCTTGAATTCTTACTTGTACAGCTCGTCCATGCCACCG |
| 5 | TACAAGCTACTTGTTCTTTTTGCAGGATCCGCCACCATGGATTGTGAGGTCAATAACGG |
| 6 | GTTCTAGAGGCTCGAGAGGCCTTGAATTCTTACTTGTTCAACCTGAGCAACCACGCATTG |

**Table S1.** Sequences of primers used in this study. Primers 1 and 2 amplified NES-PhoCl, primer 3 and 4 amplified mCherry, and primers 5 and 6 amplified NBid-PhoCl-CBid.

NES-PhoCl-linker-mCherry

atgaacctggtggacctgcagaagaagctggaggagctggagctggacgagcagcagggaggcaagcttagcgtgat
ccctgactacttcaagcagagcttccccgagggctacagctgggagcgcagcatgacctacgaggacggcggcatct
gcatcgccaccaacgacatcacaatggaggggacagcttcatcaacaagatccacttccagggcacgaacttcccc
cccaacggccccgtgatgcagaagaggaccgtgggctgggaggccagcaccgagaagatgtacgagcgcgacggcgt
gctgaagggcgacgtgaagatgaagctgctgctgaagggcggcggccactatcgcggcgactaccgcaccacctaca
aggtcaagcagaagcccgtaaagctgcccgactgccacttcgtggaccaccgcatcgagatcctgagccacgacaag
gactacaacaaggtgaagctgtacgagcacgccgtggccaagacttccaccgacagcatggacgagctgtacaaggg
tggcagcggtggcatggtgagcaagggcgaggagaccattacaagcgtgatcaagcctgacatgaagaacaagctgc
gcatggagggcaacgtgaacggccacgccttcgtgatcgagggcgagggcagcggcaagcccttcgagggcatccag
acgattgatttggaggtgaaggagggcgccccgctgcccttcgcctacgacatcctgaccaccgccttccactacgg
caaccgcgtgttcaccaagtacccacggggtaccggaggcgggggcagcggaggcgggggcagcgtgagcaagggcg
aggaggataacatggccatcatcaaggagttcatgcgcttcaaggtgcacatggagggctccgtgaacggccacgag
ttcgagatcgagggcgagggcgagggccgcccctacgagggcacccagaccgccaagctgaaggtgaccaagggtgg
cccctgcccttcgcctgggacatcctgtcccctcagttcatgtacggctccaaggcctacgtgaagcaccccgccg
acatccccgactacttgaagctgtccttccccgagggcttcaagtgggagcgcgtgatgaacttcgaggacggcggc
gtggtgaccgtgacccaggactcctccctgcaggacggcgagttcatctacaaggtgaagctgcgcggcaccaactt
ccccctccgacggccccgtaatgcagaagaagaccatgggctgggaggcctcctccgagcggatgtaccccgaggacg
gcgccctgaagggcgagatcaagcagaggctgaagctgaaggacggcggccactacgacgctgaggtcaagaccacc
tacaaggccaagaagcccgtgcagctgcccggcgcctacaacgtcaacatcaagttggacatcacctcccacaacga
ggactacaccatcgtggaacagtacgaacgcgccgagggccgccactccaccggtggcatggacgagctgtacaagt
aa

NBid-PhoCl-CBid

atggattgtgaggtcaataacggttcatctctccgagacgaatgcataacgaacttgctcgttttcggctttctgca
atcctgcagcgataattctttcagaagagaacttgacgctcttggacatgaactcccagtactcgctccacagtggg
aaggctacgatgagggatccgtgatccctgactacttcaagcagagcttccccgagggctacagctgggagcgcagc
atgacctacgaggacggcggcatctgcatcgccaccaacgacatcacaatggaggggacagcttcatcaacaagat
ccacttccagggcacgaacttcccccccaacggccccgtgatgcagaagaggaccgtgggctgggaggccagcaccg
agaagatgtacgagcgcgacggcgtgctgaagggcgacgtgaagatgaagctgctgctgaagggcggcggccactat
cgcggcgactaccgcaccacctacaaggtcaagcagaagcccgtaaagctgcccgactgccacttcgtggaccaccg
catcgagatcctgagccacgacaaggactacaacaaggtgaagctgtacgagcacgccgtggccaagacttccaccg
acagcatggacgagctgtacaagggtggcagcggtggcatggtgagcaagggcgaggagaccattacaagcgtgatc
aagcctgacatgaagaacaagctgcgcatggagggcaacgtgaacggccacgccttcgtgatcgagggcgagggcag
cggcaagcccttcgagggcatccagacgattgatttggaggtgaaggagggcgccccgctgcccttcgcctacgaca
tcctgaccaccgccttccactacggcaaccgcgtgttcaccaagtacccacggggtacccttcaaaccgatggaaac
cgatcatctcattcaaggttggggcgaattgaagcagatagtgagagtcaggaggacataatacgcaatatagctcg
acaccttgcacaggtcggtgatagcatggaccgctctatcccccccaggtttggttaacggattggccctgcaactgc
gaaacacttcaaggagtgaagaagatagaaatcgagacctggcgaccgcgcttgaacagcttcttcaagcctatccc
agagatatggagaaggaaaaaacaatgctcgtgctcgcactgctgctggcaaagaaagtagcctctaacacaccatc
cctcttgagagacgtcttccataccacggtaaatttcataaaccagaaccttaggacgtatgtgcggagtcttgctc
gcaatggtatgactgagtttcttcaacccgtgcccctcaatgcgtggttgctcaggttgaacaagtaa

**Table S2.** DNA sequences of NES-PhoCl-mCherry and NBid-PhoCl-CBid.

### Supporting Figures

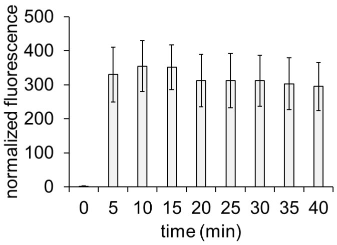


**Figure S1.** Measurement of red fluorescence from NucView 530 caspase-3 reporter in mammalian cells expressing NBid-PhoCl-CBid after irradiation with 405 nm light. Bars represent means and error bars represent standard deviation from 9 fields of view.

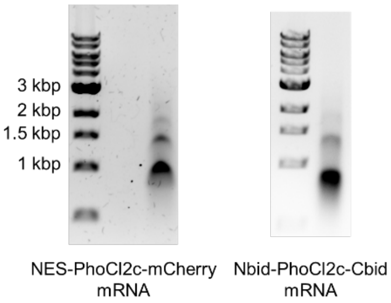


**Figure S2.** Ethidium bromide stained 0.8% agarose gel electrophoresis of mRNA generated through in vitro transcription. Tridye 1 kbp DNA ladder was included (NEB). RNA sizes do not match expected DNA ladder sizes due to differences in secondary structure in RNA, this was used for qualitative assessment of mRNA quality.

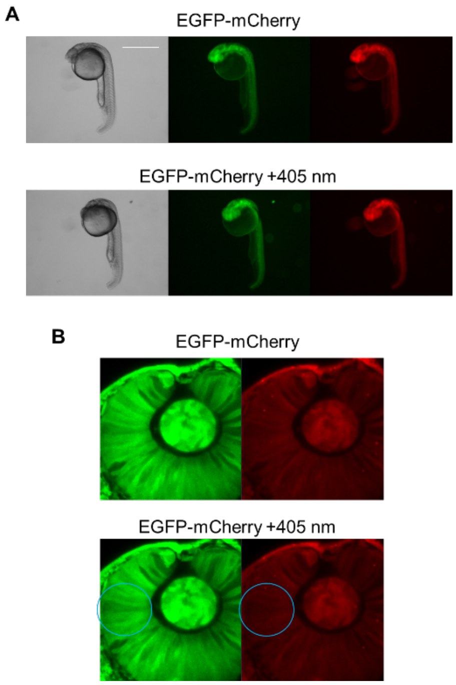


**Figure S3.** Injection of EGFP-mCherry mRNA into zebrafish embryos and irradiation with 405 nm light using the same conditions used for PhoCl experiments. A) Global irradiation with a 405 nm LED. B) Confocal 405 nm laser irradiation of the region marked by a blue circle. Scale bar = 40 μM.

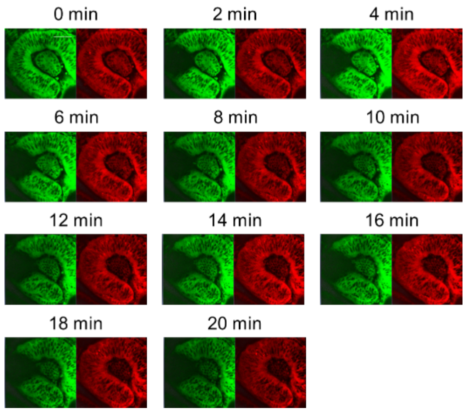


**Figure S4.** Additional images from the NES-PhoCl-mCherry timecourse experiment. Scale bar = 40 μM.

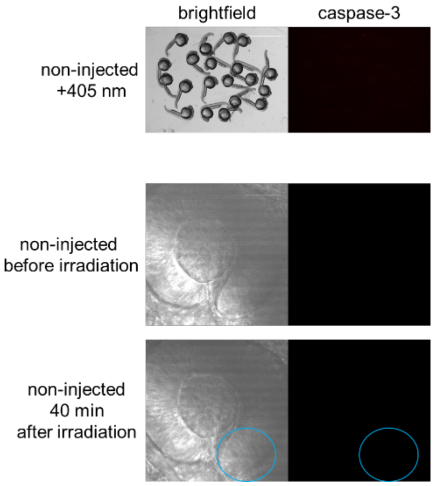


**Figure S5**. Irradiation conditions used in Nbid-PhoCl-Cbid activation experiments do not induce caspase-3 activity in non-injected embryos. Scale bars = 2 mm for the top image, 40 $\mu$M for the bottom image.

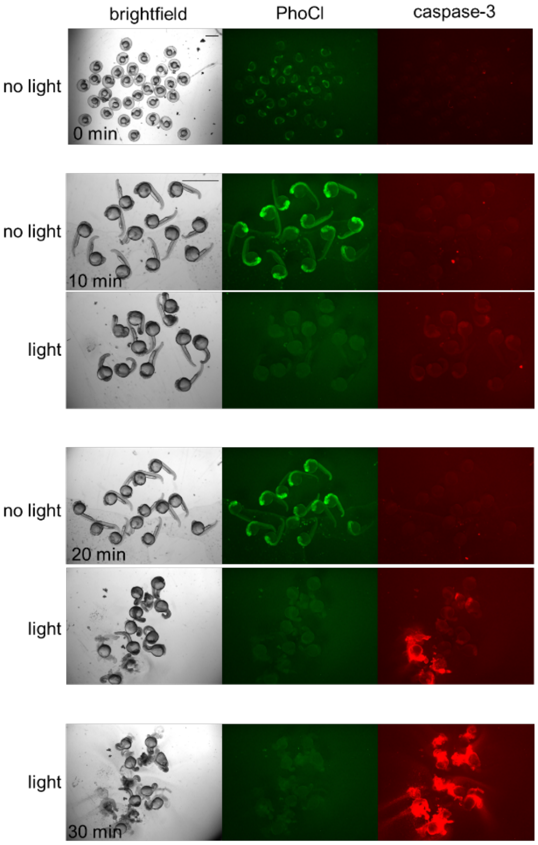


**Figure S6.** Additional images of NBid-PhoCl-CBid expressing zebrafish embryos at the designated timepoints after irradiation. Scale bar = 2 mm.

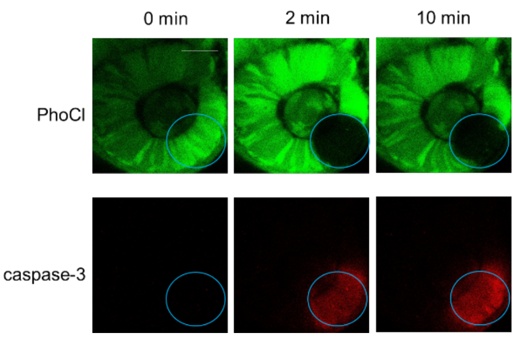


**Figure S7.** Targeted activation of apoptosis in the 24 hpf embryo eye using the confocal 405 nm laser. Scale bar = 40 μM.
